# Social costs of group life: Larger groups have less affiliative behavior and more social conflict

**DOI:** 10.64898/2026.11.10.765676

**Authors:** Brian A. Lerch, J. Kinyua Warutere, Lilian Musembei, Elizabeth A. Archie, Jenny Tung, Susan C. Alberts

## Abstract

The costs and benefits of sociality depend in part upon the size of groups. Therefore, individual behavior should vary across groups of different sizes, reflecting group size-dependent costs and benefits. Here, we test this prediction using a long-term dataset on wild female baboons (*Papio cynocephalus* x *P. anubis*). We find that individuals who live in larger groups suffer social costs: compared to animals in smaller groups, they experience more agonistic interactions, fewer grooming interactions, and sparser social networks. These social costs arise, in part, from a pattern in which relationships within dyads become more agonistic as group size increases (i.e., a given dyad exhibits fewer grooming interactions relative to agonistic interactions when it is living in a larger group). While individuals in large groups generally maintain the same number of strong bonds as individuals in small groups, they tend to develop weak bonds with a greater number of social partners that results in, on average, weaker bonds overall. Our results demonstrate that, despite potential benefits to large groups in terms of predation risk and intergroup competition, the social lives of individuals in larger groups involve less affiliation and more conflict in this wild primate population.

## Introduction

The environmental context in which social behaviors are expressed shapes their fitness consequences, and hence their evolution. Understanding the evolution of sociality thus requires understanding how the costs and benefits of social behavior vary with environmental conditions. As with any trait, the physical environment (e.g., rainfall and food availability) plays a key role in determining the costs and benefits of sociality (Cornwallis et al. 2017; Liu et al. 2020; Tilman et al. 2020; Lerch et al. 2022). However, the social environment (e.g., the identities and characteristics of an individual’s interaction partners) is also important, as it affects the expression and fitness consequences of social behavior (Akçay 2020).

Group size is an important component of the social environment that affects the evolution of social behavior (Avilés et al. 2002; Van Veelen et al. 2010; Powers et al. 2011; Peña and Nöldeke 2018; Lerch and Abbott 2020). Living in larger groups carries benefits, such as reduced predation risk, and costs, such as increased foraging effort (Van Schaik 1983; Johnsson 2003; Majolo et al. 2008; Creel et al. 2014; Lehtonen and Jaatinen 2016). Increased group size is commonly associated with increased resource competition (White 2010; Markham et al. 2015; Majer et al. 2018; Pavez-Fox et al. 2025), which often reduces survival and fecundity (Clutton-Brock et al. 1983; Lacey 2004; Kappeler and Fichtel 2012).

Group size may also influence the expression of social behaviors, affecting the group size-related costs and benefits of these behaviors. For instance, if intragroup competition increases with group size, then aggressive behavior is predicted to be more frequent in larger groups (Sirot 2000; Dubois et al. 2003); this prediction is widely supported (Saxby et al. 2010; Wheeler et al. 2013; Bailoo et al. 2018; Lu et al. 2023; Webb et al. 2025). Within groups, aggression occurs during resource competition; affiliative behavior can help mitigate the negative consequences of this conflict, facilitating group cohesion (Henzi and Barrett 1999; Radford and Du Plessis 2006; Matoba et al. 2013; Chereskin et al. 2022).

Primate social behavior is well-studied, but results concerning how it varies with group size are mixed. Some studies report that the proportion of time individuals spend grooming increases with group size, arguing that more time must be spent grooming in larger groups to maintain cohesion (Dunbar 1991; Lehmann et al. 2007). However, other meta-analyses (Majolo et al. 2008; Grueter et al. 2013) and within-population studies (Henzi et al. 1997; Sánchez-Villagra et al. 1998; Gogarten et al. 2014) fail to find a consistently positive relationship between group size and time spent grooming (but see Teichroeb et al. 2003; Jones et al. 2009). These conflicting results suggest that more work is needed to understand how both affiliative and competitive social interactions vary with group size.

We employ a comprehensive data set to understand the relationship between group size and female social behavior in a wild primate. Specifically, we analyze 150 group-years of data on female baboons in the Amboseli ecosystem of southern Kenya. In this population, affiliative social interactions predict female survival (Archie et al. 2014; Campos et al. 2020; Lange et al. 2023) and dominance rank, which is enforced by agonistic interactions, predicts female fertility (Gesquiere et al. 2018). The existence of group size effects on these interactions would therefore represent a potentially important source of variation in the expression and evolution of fitness-related behaviors. Furthermore, previous work on this population has shown that intermediate-sized groups have the most favorable range size, space use, and glucocorticoid hormone concentrations (Markham et al. 2015), and that females living in large groups have longer interbirth intervals (Gesquiere et al. 2018). Thus, baboons may experience a tradeoff between the costs of living in small groups (e.g., predation and intergroup competition) with those of living in large groups (e.g., intragroup competition). Here, we show that individuals living in large groups also suffer from a range of social costs relative to individuals living in small groups, including more agonistic interactions, fewer affiliative interactions, and weaker social bonds.

## Methods

### Study system

We use data from a long-term field study of wild baboons in the Amboseli ecosystem of Kenya, where our study subjects were adult females living in study groups from 1996-2024. This population has been under continuous observation since 1971 by the Amboseli Baboon Research Project (Alberts and Altmann 2012). All individuals in this population have both yellow baboon (*Papio cynocephalus*) and anubis baboon (*P. anubis*) ancestry as a result of recent and historical admixture (Vilgalys et al. 2022). Genetic ancestry predicts some aspects of social behavior, such as assortative mating and social partner choice (Tung et al. 2012; Franz et al. 2015; Fogel et al. 2021), but does not predict the behaviors central to this analysis (total number of grooming or agonistic interactions in which individuals engage) (McLean et al. 2023; Tuliozi et al. 2026). Further, genetic ancestry estimates are available for only a subset of females in our study. Thus, we do not consider it here. The Amboseli ecosystem is a highly seasonal semi-arid savanna: the ‘hydrological year’ in Amboseli (the ‘hydroyear’) begins on November 1^st^ (approximately the onset of the ‘short rainy season’) and ends on October 31^st^ of the following calendar year, after the annual 5-month ‘long dry season’ (Gesquiere et al. 2024).

From 1996–2024, researchers observed 307 unique adult females and 17 unique study groups, with an average of 82 adult females and 5 study groups at any given time (Southworth et al. 2026). Most females in study groups were studied since birth by observers who recognize each subject on sight and collect detailed behavioral, demographic, and developmental data during observation periods that occur 2-4 times per week per study group. We focus on adult females because they are more affected by within-group resource competition than males (Wrangham 1980; Gaulin and Sailer 1985) and because, among adults, only females are the subjects of focal sampling.

### Group size measures

We measured group size as the average number of adult females in a study group during the time period of interest. Alternative measures of group size are highly correlated with adult female group size (Supplementary Appendix S1). The average monthly number of adult females in a study group varied from 4-31, with a mean of 16.3 and a standard deviation of 6.0.

### Data collection

Data on activity budgets and social interactions were collected through two methods. First, we used 10-minute focal animal samples collected on the female study subjects (*N*=80,150 unique focal samples). Each observation day, subjects were sampled in predefined, randomized order throughout daylight hours. Two aspects of the focal sampling are most salient here. (i) The subject’s activity (feeding, resting, moving, grooming, being groomed, or other) was recorded on each minute of a 10-minute focal sample; we refer to each minute record as a ‘point’ sample. (ii) Whether the focal subject had at least one ‘near neighbor’ (i.e., a group mate of any age-sex class within 5m of the focal) was also recorded in point samples.

Second, we assigned counts of female-female grooming and agonistic behavior for each study subject using data on the 42,981 grooming interactions and 607,923 decided agonistic interactions in which they were involved. A grooming event, along with the identities of the groomer and the individual being groomed, was recorded whenever one animal systematically picked through the fur of a conspecific. A decided agonistic interaction was recorded whenever one animal (the ‘loser’) gave only submissive gestures or was displaced from its position, and another (the ‘winner’) gave aggressive or neutral, but not submissive, gestures (Hausfater 1974; Alberts et al. 2024). We classified decided agonistic interactions into two types: ‘aggressive agonistic interactions’ were those in which the winner gave exclusively aggressive gestures and the loser gave exclusively submissive gestures. ‘Nonaggressive agonistic interactions’ were those in which the winner gave neither aggressive nor submissive gestures and the loser gave only submissive gestures or was displaced by the winner (Alberts et al. 2024). Grooming and agonism data were collected throughout each observation day using ‘representative interaction sampling’: an observer moved systematically through the group while carrying out focal animal samples, simultaneously recording the identities of both participants of all grooming and agonistic interactions in their line of sight. This approach ensures a relatively even representation of all group members, with patterns of interaction frequencies and partner preferences similar to those derived from focal animal sampling. However, it yields a larger number of interactions than those that involve only the focal animal observed during focal sampling (Archie et al. 2014; Alberts et al. 2024).

### Controlling for observer effort

Calculating the relative frequency of social interactions from representative interaction data is complicated because the number of observers remains relatively constant over time, while group sizes vary. Consequently, observer effort per capita declines with increasing group size, so that more agonistic and grooming interactions per individual, and more grooming interactions per dyad, are observed in smaller groups than in larger groups. If left uncorrected, variation in observer effort affects our ability to compare relative interaction frequencies. We corrected for variation in observer effort by regressing counts of grooming interactions per dyad or agonistic interactions per group against observer effort. We calculated observer effort as the number of focal samples on adult females collected over the relevant timespan, divided by the mean number of adult females in the group during the same timespan. To measure the relative strength of dyadic social bonds, the observer effort estimate was further divided by the number of days that each dyad co-resided in the same group. We used residuals of these regressions (counts of interaction versus observer effort) as our measure of relative interaction frequency (see also Anderson et al. 2021 and Supplementary Appendix S2).

### Response variables

To investigate group size-dependent social behavior, we analyzed five sets of response variables measured at the individual, dyad, and group levels, described below.

#### Allocation of social time

We calculated the total proportion of time spent grooming, and the total proportion of time spent with at least one near neighbor (group-mate within 5m), for each adult female in each group in a given hydroyear. We counted the total number of point samples in which each of these states was recorded, and divided by the total number of point samples collected on that female in that group and hydroyear, resulting in one data point for each adult female in each hydroyear (*N*=2,604 adult female-hydroyears).

#### Relative per capita frequency of agonistic interactions

We measured the relative per capita frequency of agonistic interactions among adult females for each group-month (*N*=1,671 group-months). We counted the total number of observed agonistic interactions of each type (aggressive or non-aggressive) in a given group and month, controlling for observer effort. The <1% of months where no agonistic interactions were observed in a given group were excluded because these likely reflect sparse data collection, not a true lack of agonistic interactions.

#### Proportion of positive interactions per dyad

To understand how relationships within specific dyads vary with group size, we examined the proportion of positive interactions within each adult female dyad during each group-hydroyear. For each adult female dyad co-resident in a given group and hydroyear, we counted the number of times the dyad was observed grooming and the number of times the dyad was observed engaging in agonistic interactions. We then used these counts to model the probability any given interaction for a dyad over the course of a hydroyear was grooming rather than agonistic. We refer to dyads for which this probability is higher as having a more positive ‘relationship tenor’. We restricted our analysis to dyads that interacted at least 6 times during the hydroyear (*N*=9,937 dyad-hydroyears).

#### Social bonding metrics

We calculated average values of the number of social partners and bond strength between partners for each group-hydroyear. For number of social partners, we calculated the number of other adult females with whom each adult female in a group groomed at least once in the given hydroyear. For bond strength, we calculated the residuals from the regression of the number of grooming interactions against observer effort for every female-female dyad that groomed at least once in a given group-hydroyear (*N*=150 group-hydroyears). We then calculated means for each group-hydroyear for each measure. To improve model fit, we transformed the initial estimates of bond strength, which included negative values, to have a minimum of 1, which we refer to as ‘shifted bond strength’. We repeated both analyses with a restricted dataset that counted social partners only if they had bonds that were stronger than the average bond across all group-hydroyears (i.e., those having a positive residual given observer effort; ∼50% of all bonds).

#### Social network metrics

We computed three metrics of grooming-based adult-female social networks for each group and hydroyear (*N*=150 group-hydroyears). We assumed that a dyad was connected in the network if they had a stronger grooming bond than the average bond (controlling for observer effort) between any two adult females observed to groom at least once across all group-hydroyears. Our social network metrics assess different aspects of adult female social network connectedness. (i) The clustering coefficient measures whether the group consists of tighly connected subgroups and is the proportion of all triads in a given group-hydroyear that are “closed” (i.e., the proportion of triads where all three dyads have above-average bond strength, and where triads are sets of three females with at least two dyads having above-average bond strength). (ii) The network density measures overall group connectedness and is the proportion of dyads in the network that have an above-average bond strength, relative to the maximum number of possible dyads in the network. (iii) The diameter measures whether the group is well connected in terms of some individuals being separated by many steps (high diameter means poorly connected) and is calculated by measuring the shortest path between each dyad in the network and then identifying the longest of these paths.

### Statistical procedure

We investigated how each response variable varied with group size using generalized linear mixed effect models. For each response variable, we used the glmmTMB package (Brooks et al. 2017) to fit two models, one with only a linear fixed effect of group size and one with both a linear and a quadratic fixed effect of group size. Each model included random effects of group identity and individual/dyad identity (when applicable). Model assumptions were tested using the DHARMa package (Hartig 2022). If model diagnostics revealed a significant deviation from the assumed distribution, we removed a random effect as needed for a better match to assumptions. We accounted for overdispersion by including an observation-level random effect where each observation was assigned a unique random intercept drawn from a normal distribution with a mean of zero and an estimated variance (Harrison 2014). We used AIC to identify the best-supported model. Statistical analyses were conducted in R (R Core Team 2022) with visualization in Mathematica (Wolfram Research Inc. 2022).

In analyzing the proportion of positive interactions per dyad, we sought to differentiate two alternative explanations for a potential decline in the proportion of positive interactions with group size. Such a decline could result from a within-dyad effect, in which the proportion of positive (grooming) compared to negative (agonistic) interactions for specific dyads (i.e., their relationship tenor) was lower in larger groups. Alternatively, females might interact with different subsets of groupmates in groups of different sizes. For instance, if two females lack a bond and avoid each other in small groups, but engage in increasing rates of agonistic (but not affiliative) interactions as group size increases, we would also observe a higher relative frequency of agonistic interactions without a change in the tenor of specific relationships. To differentiate these possibilities, we modeled the proportion of positive interactions per dyad as a function of the mean group size experienced by each dyad *N̅_ij_* (which differs between dyads) and the deviation in group size a dyad experienced in year *t* from their mean value, (the dyad’s group size anomaly; *N_ijt_* − *N̅_ij_*) (which differs within-dyads). We included dyad identity as a random effect. A significant effect of deviation in group size within dyads indicates that the tenor of particular dyadic relationships changed with group size.

## Results

### Females in larger groups spent less time grooming and less time with a near neighbor

Females in larger groups spent less time grooming (*p* < 10^−12^; Table 1; Fig. 1a; Table S3.1). Females in the smallest groups (∼4 adult females) spent 7.1% of their time grooming while females in the largest groups (∼30 adult females) spent 3.0% of their time grooming, a ∼60% difference.

**Figure 1.**
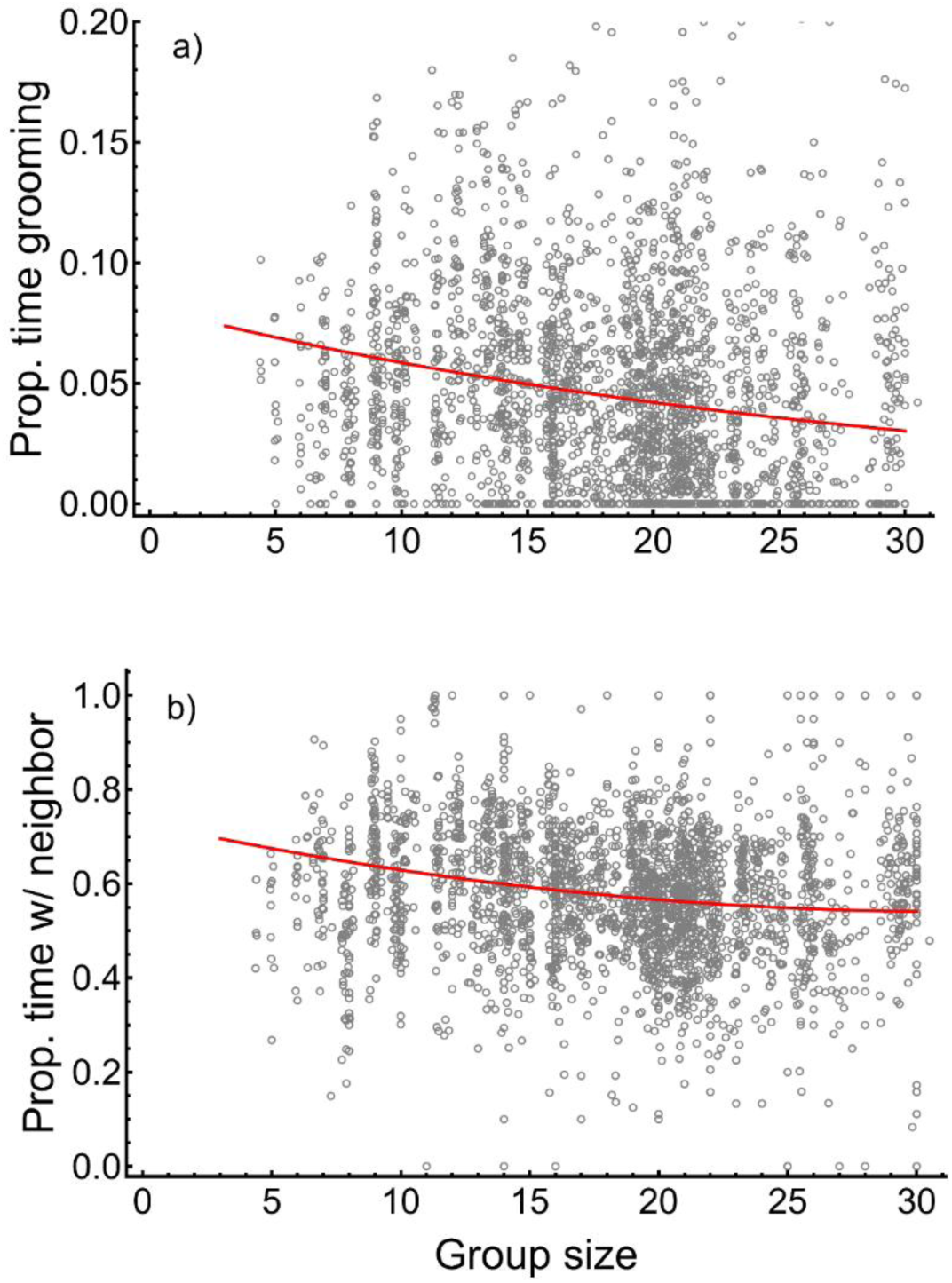
Adult females spent the least time (a) grooming and (b) with a near neighbor in the largest groups. Each circle represents the time allocation of one female over one hydroyear and the red lines show the best-supported models.

**Table 1.** Summary of results from the best-supported model for each primary research question involving the 307 adult female sujects (Int = intercept; *x* = group size; *x̅* = mean group size experienced by a dyad; *T* = total number of dyadic interactions).

| Response category | Response variable | Unit of analysis and sample size | Best-supported model | p-values | Result |
|---|---|---|---|---|---|
| Allocation of social time | Proportion of time adult females spend grooming | Female-hydroyear (*N* = 2,604) | log(*y*) = −2.508 − 0.033*x* | Int: < 10<sup>−15</sup><br>*x*: < 10<sup>−12</sup> | Females spend less time grooming in large groups |
|  | Proportion of time adult females spend with near neighbor | Female-hydroyear (*N* = 2,604) | *y* = 0.730 − 0.012*x* + 0.0002*x*<sup>2</sup> | Int: < 10<sup>−15</sup><br>*x*: < 10<sup>−3</sup><br>*x*<sup>2</sup>: 0.032 | Females spend less time with neighbors in large groups; this effect attenuates in the largest groups |
| Agonistic interactions | Relative monthly per capita frequency of aggressive agonistic interactions (residual) for adult females | Group-month (*N* = 1,671) | *y* = −0.380 + 0.029*x* | Int: < 10<sup>−4</sup><br>*x*: < 10<sup>−13</sup> | Females engage in more aggressive agonistic interactions in large groups |
|  | Relative monthly per capita frequency of non-aggressive agonistic interactions (residual) for adult females | Group-month (*N* = 1,669) | *y* = −0.835 + 0.070*x* − 0.0009*x*<sup>2</sup> | Int:< 10<sup>−8</sup><br>*x*: < 10<sup>−6</sup><br>*x*<sup>2</sup>: 0.019 | Females engage in more non-aggressive agonistic interactions in large groups |
| Proportion positive interactions | Proportion of interactions within adult female dyads that were positive (grooming) vs. negative (agonistic) | Dyad-hydroyear (*N* = 9,937) | *logit*(*y*) = −0.625 − 0.022(*x* − *x̄*) − 0.033*x̄* + 0.007*T* | Int: < 10<sup>−4</sup><br>*x* − *x̄*: < 10<sup>−9</sup><br>*x̄*: < 10<sup>−5</sup><br>*T*: < 10<sup>−15</sup> | Dyads have the highest proportion of positive interactions in small groups |
| Social bonding metrics | Average strength of all bonds among adult females | Group-hydroyear (*N* = 150) | log(*y*) = 1.936 − 0.134*x* + 0.003*x*<sup>2</sup> | Int: < 10<sup>−15</sup><br>*x*: < 10<sup>−9</sup><br>*x*<sup>2</sup>: < 10<sup>−7</sup> | Mean female bond strength is greater in small groups, especially so in the smallest groups |
|  | Average strength of strong bonds among adult females | Group-hydroyear (*N* = 150) | log(*y*) = 1.032 − 0.083*x* + 0.002*x*<sup>2</sup> | Int: < 10<sup>−12</sup><br>*x*: < 10<sup>−5</sup><br>*x*<sup>2</sup>: 0.001 | Mean female strong bond strength is greater in small groups, especially so in the smallest groups |
|  | Average number of adult female social partners for adult females | Group-hydroyear (*N* = 150) | *y* = −1.731 + 1.191*x* − 0.028*x*<sup>2</sup> | Int: 0.199<br>*x*: < 10<sup>−12</sup><br>*x*<sup>2</sup>: < 10<sup>−8</sup> | Females in large groups have more social partners on average; this effect attenuates in the largest groups |
|  | Average number of social partners with whom adult females had a strong bond | Group-hydroyear (*N* = 150) | *y* = 4.689 | N/A | The average number of social partners connected by strong bonds is independent of group size |
| Social network metrics | Clustering coefficient in the adult female social network | Group-hydroyear (*N* = 150) | *y* = 1.084 − 0.070*x* + 0.001*x*<sup>2</sup> | Int: < 10<sup>−15</sup><br>*x*: < 10<sup>−9</sup><br>*x*<sup>2</sup>: < 10<sup>−5</sup> | Female social networks are more clustered in small groups, especially so in the smallest groups |
|  | Density of the adult female social network | Group-hydroyear (*N* = 150) | *y* = 1.138 − 0.080*x* + 0.002*x*<sup>2</sup> | Int: < 10<sup>−15</sup><br>*x*: < 10<sup>−15</sup><br>*x*<sup>2</sup>: < 10<sup>−8</sup> | Female social networks are denser in small groups, especially so in the smallest groups |
|  | Diameter of the adult female social network | Group-hydroyear (*N* = 150) | *y* = 4.575 | N/A | Social network diameter is independent of group size |

Adult females in the largest groups also spent ∼20% less time with a neighbor within 5m than females in the smallest groups (*p* < 10^−3^; Table 1; Fig. 1b Table S3.1). The best-supported model included both a negative linear and positive quadratic effect of group size, indicating that the time spent with a near neighbor decreased more sharply from small to medium-sized groups than it did from medium-sized to large groups.

### Per capita agonistic interactions were more frequent in larger groups

The relative frequency of aggressive and non-aggressive agonistic interactions among adult females increased with group size. The best-supported model for aggressive agonistic interactions predicted a positive linear increase with group size (*p* < 10^-13^; Table 1; Fig. 2a). The best-supported model for non-aggressive agonistic interactions predicted a non-linear increase with group size, with the rate of increase gradually slowing in the largest groups (*p* = 0.019, Table 1; Fig. 2b).

**Figure 2.**
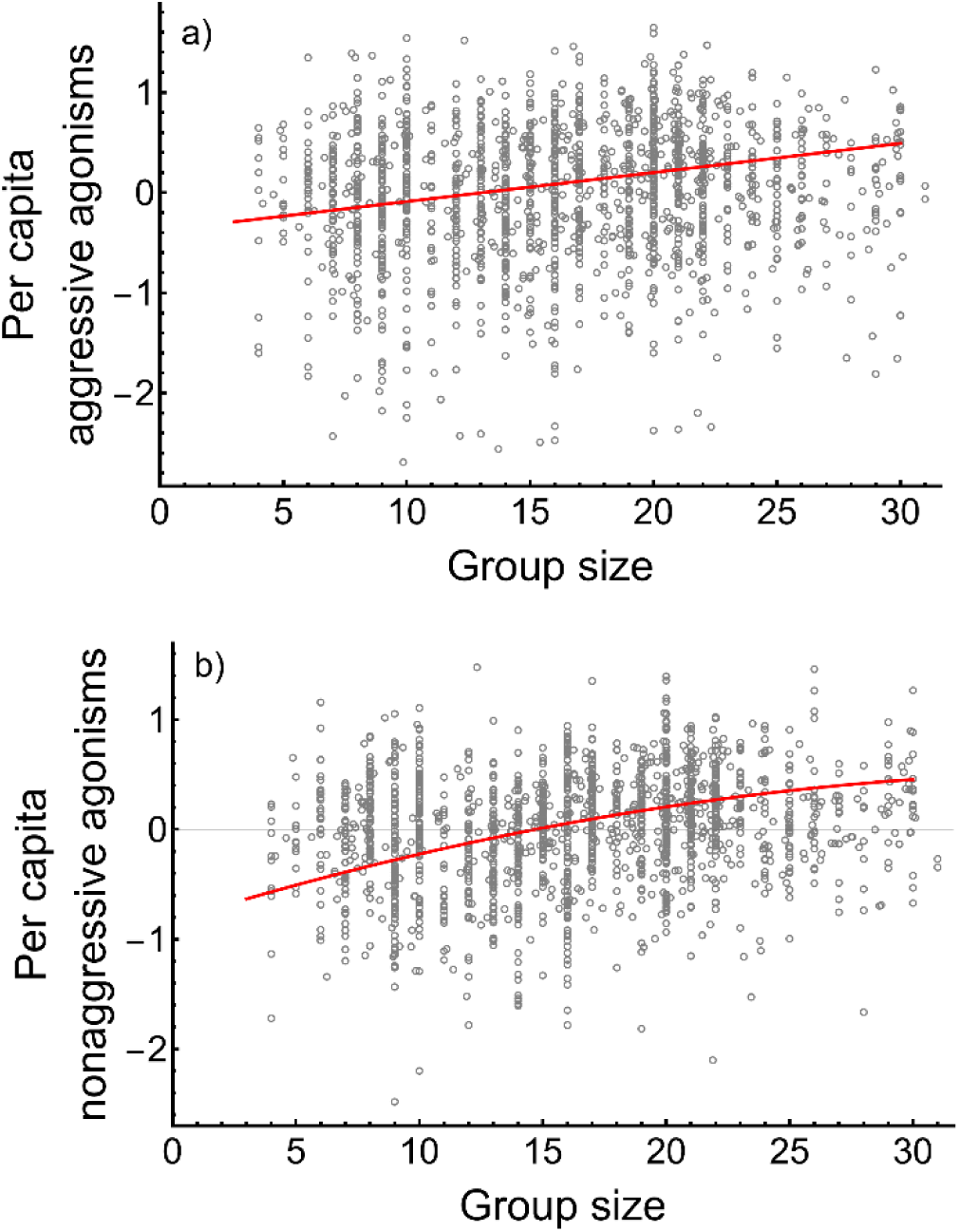
The relative per capita frequency of (a) aggressive and (b) nonaggressive agonistic interactions involving two adult females is higher in larger groups (plotted as the residual of agonism counts, controlling for observer effort; Supplementary Appendix S2). Each circle represents the number of per-capita aggressive agonistic interactions among adult females in one social group in one month, controlled for observer effort. The red line is the best-supported model.

### Dyads were more agonistic in larger groups

Within dyads, relationships tended to have more negative relative to positive interactions (i.e., they had a more negative tenor) as the size of their group increased. Specifically, increasing group size by one adult female was associated with an approximately 0.4% decrease in the proportion of positive interactions within a given dyad (a lower proportion of interactions were grooming compared to agonistic; *p* < 10^-9^; Table 1; Fig. 3; Table S5.1). This effect held even taking into account variation between dyads, though dyads in larger groups had a more negative tenor overall (*p* < 10^-5^; Table S5.1). The proportion of positive interactions nearly halved in the largest group relative to the smallest. This suggests that dyads not only become more agonistic in larger groups, but also that females that do not interact in small groups begin to interact agonistically as their group grows.

**Figure 3.**
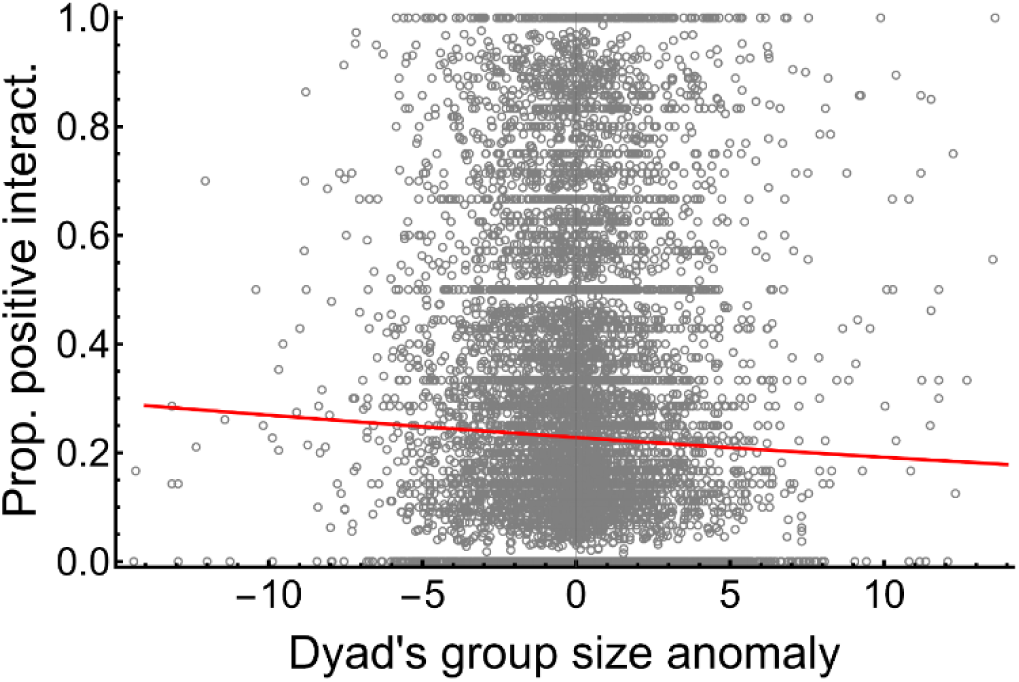
The proportion of positive interactions within adult female dyads is higher (relationship tenor is more positive) when group size is smaller than the average group size experienced by those dyads (i.e., when the group size anomaly is negative). Each circle represents one adult female dyad in one hydroyear. The vertical axis is the proportion of all interactions over the course of a hydroyear that are grooming (positive) rather than agonistic (negative) interactions for that dyad. The red line is the best-supported model.

To further explain variation in relationship tenor, we included a measure of the total number of interactions in the model. Dyads that interacted more frequently tended to have a higher proportion of positive interactions (*p* < 10^−15^; Table S5.1). The proportion of positive interactions was ∼3x higher in dyads that interacted the most compared to those that interacted the least.

### Females in larger groups had more social partners, but weaker bonds

We found a non-linear effect of group size on mean bond strength: mean bond strength per group-hydroyear declined as group size increased from small to medium but did not decrease substantially as group size increased from medium to large. This pattern was consistent whether we considered bonds of any strength or only those that were stronger than the mean across all group-hydroyears (*p* < 0.001; Table 1; Fig. 4ab; Table S6.1).

Mean bond strength may be greater in small groups compared to larger groups if individuals increase their total number of social partners in larger groups at the expense of individual bonds. Indeed, females had more total social partners in medium and large groups compared to small groups. The best model included both linear and quadratic group size effects: in groups with up to ∼22 adult females (one standard deviation above the mean), each additional adult female predicted ∼0.5 additional social partners per female (*p* < 10^−8^; Table 1; Fig. 4c; Table S6.2). Group size did not predict the mean number of social partners with strong bonds (*p* = 0.292; Table 1; Fig. 4d; Table S6.2).

**Figure 4.**
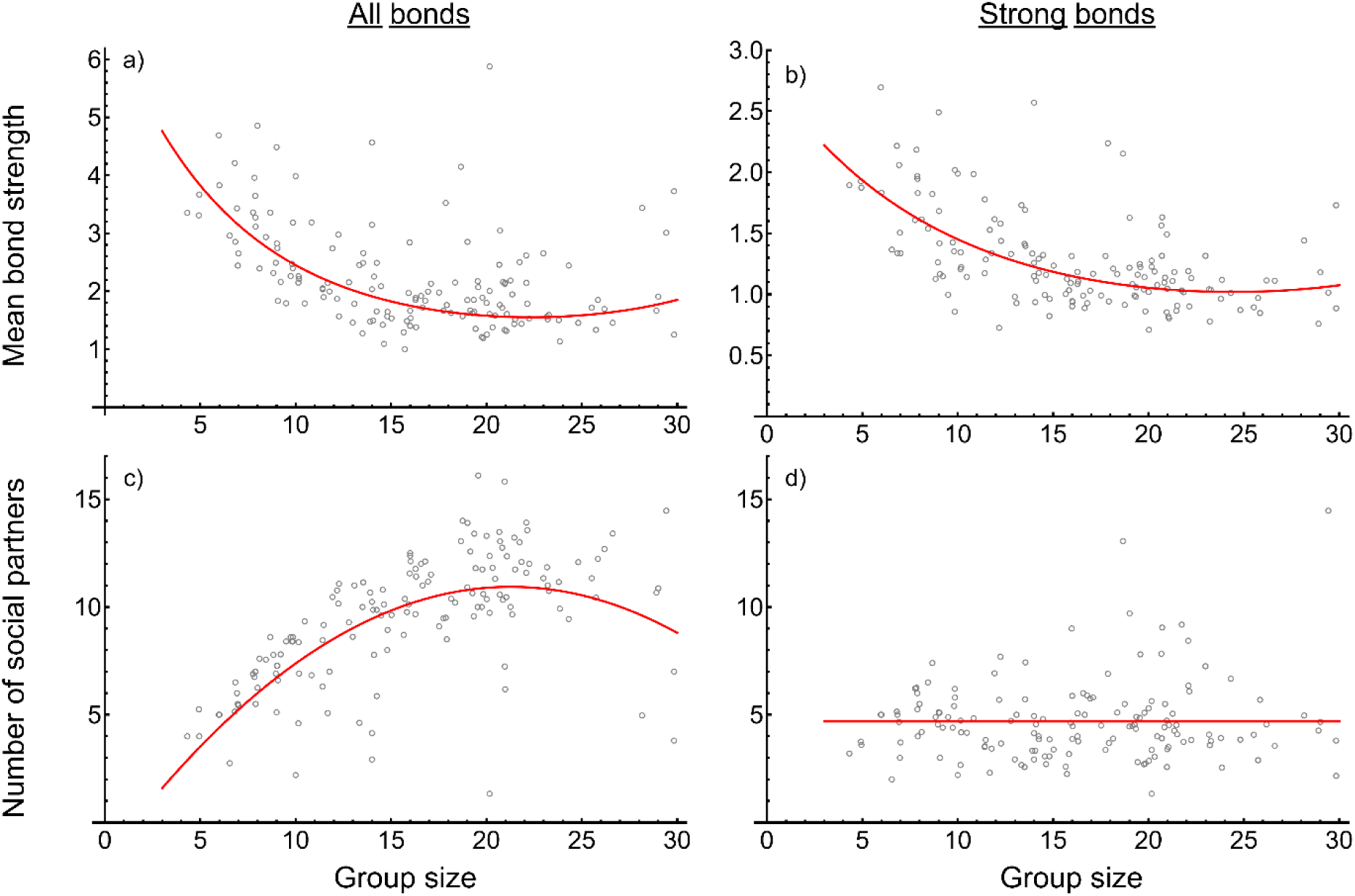
Larger groups have weaker measures of social bonding. (a) All bonds between adult females and (b) strong bonds between adult females were weaker in larger groups. (c) The mean number of total social partners for adult females tended to be larger in groups with more adult females. (d) The mean number of social partners with whom a female had strong bonds did not change with group size. Each circle represents one social group in one hydroyear and shows mean strength of bonds in a given group-hydroyear (a, b) or the mean number of social partners in a given group-hydroyear (c, d) either including all bonds (a, c) or restricted to only bonds of above average strength (b, d). In panel (a) bond strength was shifted such that the weakest bond was set to 1 to meet model assumptions (see Methods), meaning the vertical axis scales are not comparable between (a) and (b). Red lines show best-supported models.

Females in larger groups had more, but weaker, bonds than females in smaller groups. To assess whether the total social effort of females changed in groups of different size, we analyzed the mean weighted degree for each group-hydroyear (i.e., the sum of all of bond strengths for each adult female). The mean weighted degree was significantly higher in larger groups when considering all bonds, but significantly lower in larger groups when restricting to bonds of above average strength (*p* < 0.01; Table S6.1; Fig. S6.1).

### Social networks were less cohesive in larger groups

The clustering coefficient (proportion of triads that are closed; *p* < 10^−5^; Table 1; Fig. 5a; Table S7.1) and the density of social networks (proportion of possible bonds that exist; *p* < 10^−8^; Table 1; Fig. 5b; Table S7.1) were lower in larger groups. The social networks of larger groups were therefore less clustered and more weakly connected. The best-supported models included both a negative linear and positive quadratic group size effect, indicating that the decline was particularly steep in the transition from small to medium-sized groups. As expected given that our measure of bond strength which was used to create social networks was controlled for observer effort, these results were insensitive to introducing additional controls for observer effort (Table S7.2). The diameter of the social networks was not significantly related to group size, though there was a trend towards larger diameters in larger groups (*p* = 0.082; Table 1; Fig. 5c; Table S7.1).

**Figure 5.**
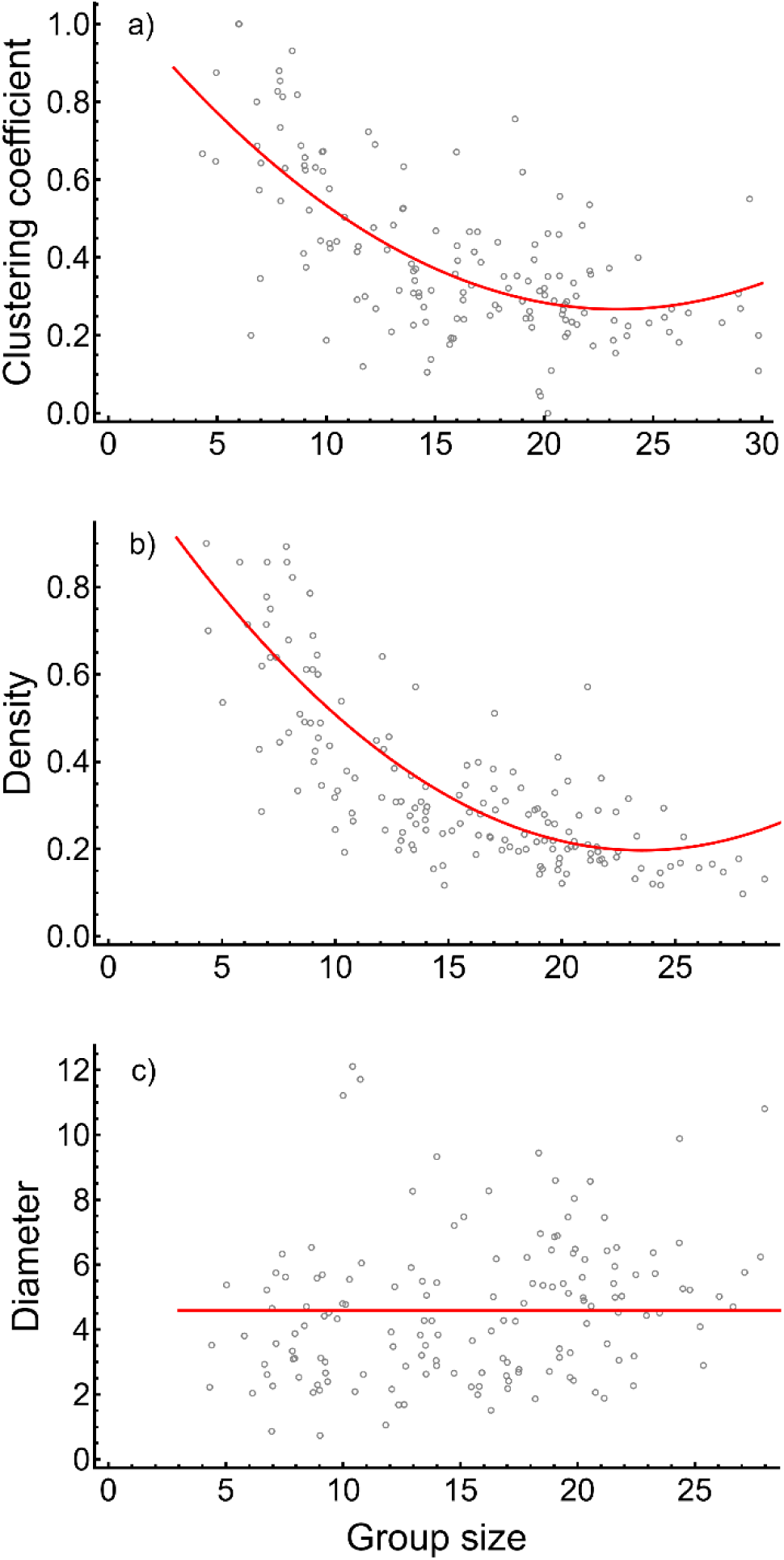
Social networks were more sparsely connected in large groups. The (a) clustering coefficient and (b) density of the adult female social network were lower in larger groups. (c) Group size did not predict the diameter of the social network. Each circle represents one social group in one hydroyear and shows one metric of a social network for a given group-hydroyear. The red lines are the best-supported models.

## Discussion

Our results suggest that social costs increase with group size in baboons. In larger groups, individuals spent less time grooming and agonistic interactions were relatively more frequent per capita than in small groups. Additionally, the tenor of dyadic relationships—i.e., the proportion of interactions within the dyad involving grooming versus agonisms—was increasingly weighted towards agonisms in larger groups. While females had the same number of strong social bonds in large groups as in small groups, they had a greater total number of social partners and weaker bonds overall, a result also found in chacma baboons (*P. ursinus*) (Silk, Seyfarth, and Cheney 2018). These individual-level behavioral differences had group-level consequences: social networks were less well-connected in large groups.

The largest primate groups are thought to be under the greatest energetic stress (Stacey 1986; Majolo et al. 2008; Markham et al. 2015; Markham and Gesquiere 2017). We suspect that increased within-group competition in large groups imposes net energetic costs for Amboseli baboons, and that this underlies the observed social costs. Large groups in Amboseli spend more time foraging than all other groups, travel greater distances each day, and have members with higher concentrations of glucocorticoid metabolites in fecal samples than members of intermediate-sized groups (Markham et al. 2015). Females in larger groups also have longer periods of lactational amenorrhea, longer interbirth intervals, and somewhat lower thyroid hormone concentrations than females in smaller groups (Hill 2000; Altmann and Alberts 2003; Gesquiere et al. 2024, 2018).

Large group size could confer benefits that we did not measure. Living in larger groups has been associated with diverse benefits, including improved foraging efficiency and thermoregulation (Villegas et al. 2013; MacNulty et al. 2014; Snijders et al. 2021). In baboons, two likely benefits of large group size are protection against predation and higher success in intergroup competition. Predation risk declines with group size in many animals, including several primates (Hill and Lee 1998; Wrona 1991; Bettridge and Dunbar 2013; Beauchamp et al. 2021). Furthermore, large groups in Amboseli perform better in between-group competition, allowing their members to have improved access to resources compared to small groups (Markham et al. 2012, 2015; Markham and Gesquiere 2017).

For these reasons, past studies in the Amboseli baboons suggest intermediate-sized groups are optimal (Markham et al. 2015). However, most of the models we present here indicate a monotonically negative effect of group size. While social costs declined slightly in the largest groups (e.g., bond strength and social network metrics, Figs 4, 5), this likely reflects the challenges of measuring of dyadic relationships at the largest group sizes. Specifically, our control for observer effort only applied to dyads that were observed grooming, but any given dyad is less likely to be seen grooming in the largest groups, possibly creating a spurious non-monotonic relationship at large group sizes. The potential benefits to living in large groups for Amboseli baboons thus represents an area for future study.

How can primates regulate group size to balance its costs and benefits? Permanent group fission can reduce group size and, presumably, the costs of within-group competition (Dittus 1988; Barton et al. 1996; Ron 1996; Feder et al. 2025). Fissions in blue monkeys (*Cercopithecus mitis*) have clear fitness benefits consistent with reduced within-group competition (e.g., improved diet, less agonism, and improved demographic rates) (Wakeford and Cords 2025a). However, fissions in Amboseli are rare, occurring approximately once per adult female lifetime (every ∼20 years; Van Horn et al. 2007; Southworth et al. 2026). Thus, fissioning may be limited by other considerations. For instance, fissions could produce groups that are too small, resulting in elevated mortality and vulnerability in between-group competition (Isbell et al. 1991; Markham et al. 2015; Majolo et al. 2020). After fissions, groups are unlikely to persist at the optimal size for long because optimally sized groups, by definition, have the highest birth rate and/or the lowest death rate, and therefore grow the fastest. Thus, the benefits of optimizing group size are ephemeral and optimal group sizes will rarely be observed (Sibly 1983; Giraldeau 1988; Higashi and Yamamura 1993; Halloway et al. 2020). Finally, fissions may impose social costs. While each animal benefits from remaining with its closest social partners, animals in groups with complex social networks cannot easily do so during a fission (Lerch et al. 2021). Indeed, even females who succeed in maintaining some strong social bonds post-fission, lose some strong bonds as well (Lerch et al. 2021; Wakeford and Cords 2025b).

Given these constraints on fissions, the apparent net energetic costs of living in large groups may sometimes be unavoidable. Reduced time spent socializing and increased aggression in large groups may be individuals’ best strategy to meet energetic demands (Dunbar 1992; Dunbar 2025). Similarly, increased aggression may improve access to resources within a competitive environment. Thus, these “social costs” may be females making the best of a bad situation, and represent adaptive plasticity. Animals may differ in their ability to adjust to large groups. For example, age or status may lead to systematic differences in plastic responses to group size, as in blue monkeys, where high-ranking females are responsible for most of the increased aggression in large groups (Klass and Cords 2015). In addition, individuals may exhibit heritable differences in behavior that could affect the extent of behavioral plasticity (e.g., Wilson et al. 2009; Brent et al. 2013; Yoshida et al. 2016; Araya-Ajoy and Dingemanse 2017).

Social plasticity in response to changes in intragroup competition can play out in several ways. For example, captive primates living at high density often do not become more aggressive in spite of increased opportunities for conflict, possibly because they decrease their rates of all social interactions, including positive ones, to make conflict less likely (Judge and de Waal 1993; Aureli et al. 1995; Videan and Fritz 2007). However, in the Amboseli baboons and in previous meta-analyses of wild populations (Wheeler et al. 2013; Cowl and Shultz 2017), agonistic interactions become relatively more frequent in larger groups. This captive-wild difference is most likely explained by the ability of primates in natural populations to spend less time with near neighbors when groups are large, decoupling group size from spatial density (Fig. 1b; Chancellor and Isbell 2009).

We see two important future directions. First, while within-group competition likely drives the social costs of living in large groups, we cannot confirm a causal effect of group size on behavior in this observational study. An alternative explanation lies in ecosystem-level changes in Amboseli and population growth in the Amboseli baboons. Specifically, the ecosystem has shifted from a semi-arid wooded savanna toward a more open habitat in the last few decades (Western 2007). Anthropogenic effects, driven by increased human and livestock density, have altered vegetation composition, hydrology, and herbivore grazing patterns in Amboseli (Okello and D’Amour 2008; Western et al. 2021; Western and Mose 2021). At the same time, the baboon population has been growing as it recovers from a major population crash of unknown origin in the 1960s (Altmann et al. 1985; Samuels and Altmann 1991; Storz et al. 2002). Since groups in this population typically grow until they fission, temporal changes in the ecosystem and population size correlate with group size, potentially confounding our results. The importance of ecosystem-level effects on baboon demography and behavior represents a key future direction. Second, it remains unclear whether living in large groups entails net fitness benefits or costs in this population. Individuals in larger groups have more social partners (which carries survival benefits; Archie et al. 2014), but weaker social bonds (which carries survival costs; Campos et al. 2020; Lange et al. 2023). Future work assessing the relative importance of relationship quantity versus quality on fitness will help to disentangle these effects.

