## Supplement for "Social costs of group life: Larger groups have less affiliative behavior and more social conflict"

### Supplementary Appendix S1: Measures of group size

In the main text, we measured group size as the number of adult females, since our primary analysis focused on interactions between adult females. The average number of adult females per month in the study groups within this data set was 16.3 ± 6.0 (S.D.) individuals (range 4.0-31.0 Fig. S1.1a). Measuring only adult female group size is unlikely to influence our results because potential alternative measures of group size are highly correlated with the number of adult females (e.g., $r^2$ between total group size and adult female group size is 0.86; Fig. S1.1b).

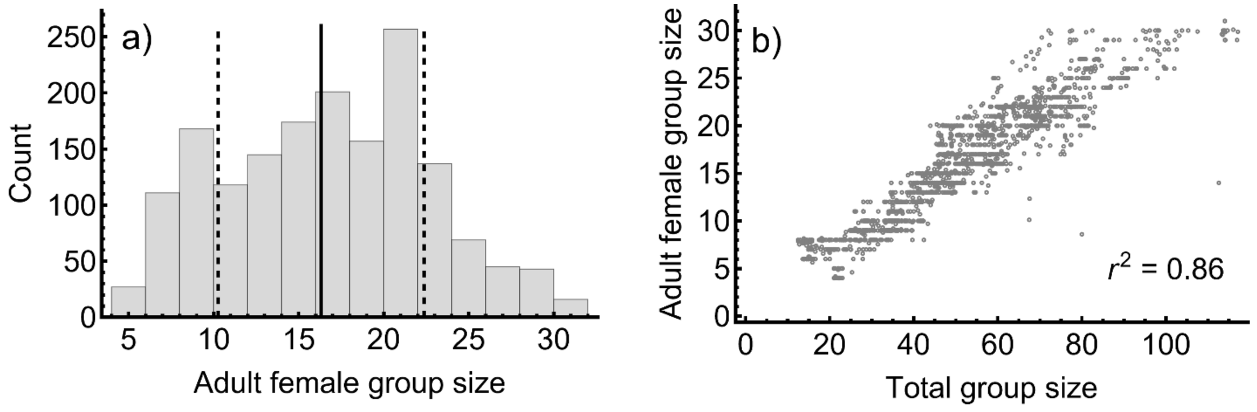


**Figure S1.1.** (a) Frequency distribution of average monthly number of adult females. Solid vertical line indicates the mean and dashed vertical lines indicate one standard deviation from the mean. (b) Adult female group size is tightly correlated with total group size.

**Supplementary Appendix S2: Controlling for observer effort**

Controlling for observer effort for each social group (calculated based on the number of focal samples per adult female in the social group) is essential in order to compare the relative frequencies of social interactions between groups of different sizes. For grooming data, this process was automated using the ramboseli R package (Campos 2021) and is explained in detail in Anderson et al. (2021). A procedure for controlling for observer effort in agonism data has not been similarly automated, so we explain our approach here using aggressive agonistic interactions between adult females as an example. We used a parallel procedure for nonaggressive agonistic interactions.

In our approach, controlling for observer effort requires three quantities: the intended response variable (in our example, per capita number of agonistic interactions in a group in a given month), the number of focal samples carried out on the group in a given month, and the average group size in the group in a given month (measured as the number of adult females in the group in that month). For measures of social bonds, controlling for observer effort also requires the number of days that a given dyad was co-resident in the same group in a given month (the actual number of days both members of dyad were inferred to live a given group, not just days they were observed).

Observer effort is defined as the number of focal samples collected on adult females divided by the group size in a given month. In the case of social bonds, this quotient is then divided by the number of co-resident days. Next, we regress the log of the per capita number of agonistic interactions against the log of observer effort, which reveals a strong positive correlation between observer effort and the number of per capita agonistic interactions observed (see Fig. S2.1 for the example of aggressive agonistic interactions between adult females). In other words, the per capita number of agonistic interactions that were observed is confounded by the fact that the number of observers remained relatively constant over time, while social group sizes varied.

To address this confound, we take advantage of the fact that the regression of agonistic interactions against observer effort provides an expected number of agonistic interactions for a group, given a certain amount of observer effort. Thus, points that fall above the regression line in Fig. S2.1 represent cases where observers recorded more agonistic interactions than expected given observer effort and points that fall below the regression line represent cases where observers recorded fewer agonistic interactions than expected given observer effort. We consider the residuals from this regression to represent a biologically valid measure of the relative frequency of per capita agonistic interactions, and we use these residuals as response variables in statistical analyses of the number of per capita agonistic interactions (e.g., Fig. 2).

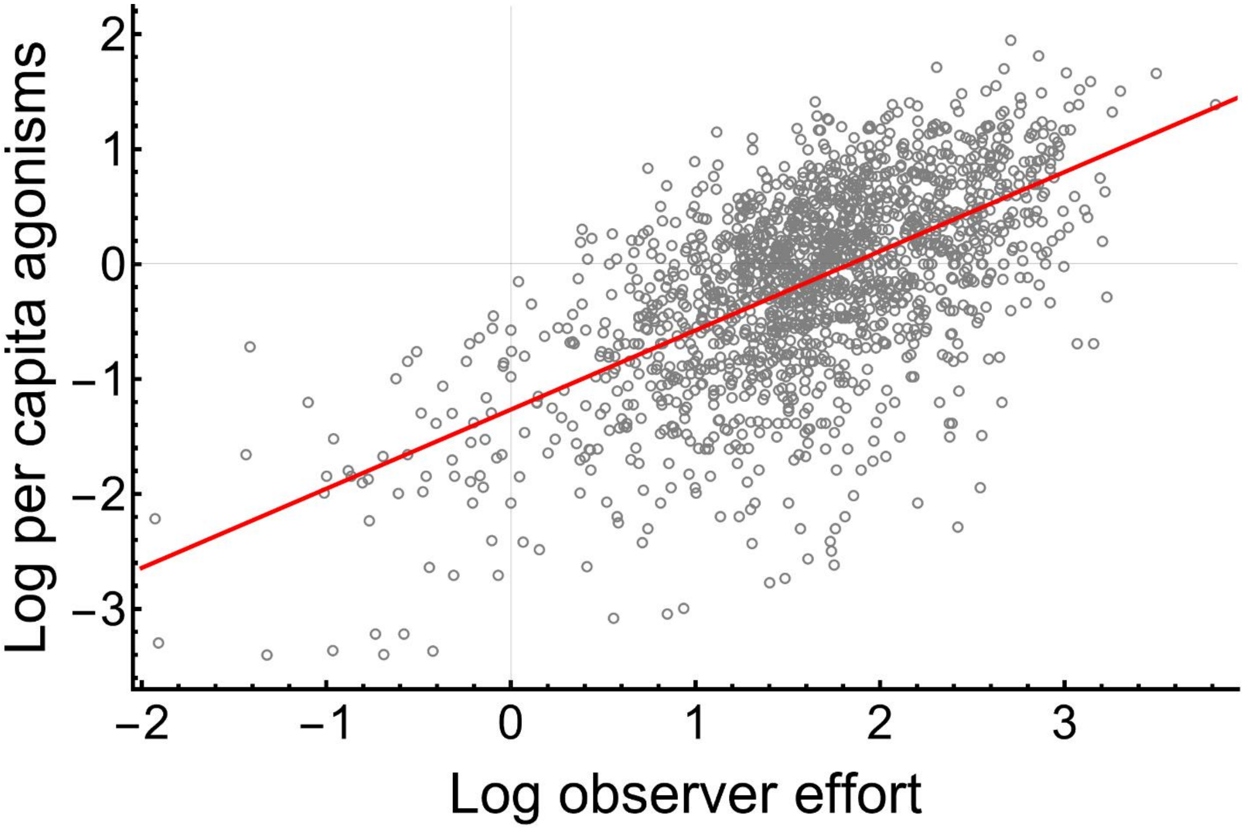


**Figure S2.1**. More non-aggressive agonistic interactions were observed between any two adult females in groups with more observer effort ($y = -1.266 + 0.689x$; $p \ll 0.001$). Circles show the log of relative per capita frequency of aggressive agonistic interactions between adult females in the group, against the log of observer effort for a given group averaged over one month.

**Supplementary Appendix S3: Detailed statistical methods and results for time spent grooming and with a near neighbor**

**Table S3.1**. Detailed model results for time spent grooming and with a near neighbor with response variables shown in italics ($x$ = group size). AIC difference from best-supported model provided for each response variable. P-values associated with the model fit are listed in the third to last column (Int = intercept). All intercepts were statistically significantly different from zero; other statistically significant p-values are bolded. Response distribution and random effects included in each model are noted in the final two columns. See Fig. S3.1 for raw distribution of proportion of time grooming, which includes many individuals that never groom. A summary of the best-supported model for each response variable can be seen in Table 1.

| **Form** | **Model** | **ΔAIC** | **p-values** | **Distribution** | **Random effects** |
|---|---|---|---|---|---|
| | *Proportion time grooming* | | | | |
| Linear | $\log(y) = -2.508 - 0.033x$ | 0 | Int: $< 10^{-15}$<br>$\boldsymbol{x: < 10^{-12}}$ | Tweedie | Group id<br>Individual id |
| Linear + quadratic | $\log(y) = -2.413 - 0.045x + 0.0003x^2$ | 1.8 | Int: $< 10^{-15}$<br>$x$: 0.068<br>$x^2$: 0.635 | Tweedie | Group id<br>Individual id |
| | *Proportion time with near neighbor* | | | | |
| Linear + quadratic | $y = 0.730 - 0.012x + 0.0002x^2$ | 0 | Int: $< 10^{-15}$<br>$\boldsymbol{x: < 10^{-3}}$<br>$\boldsymbol{x^2: 0.032}$ | Gaussian | Group id<br>Individual id |
| Linear | $y = 0.671 - 0.005x$ | 2.6 | Int: $< 10^{-15}$<br>$\boldsymbol{x: < 10^{-13}}$ | Gaussian | Group id<br>Individual id |

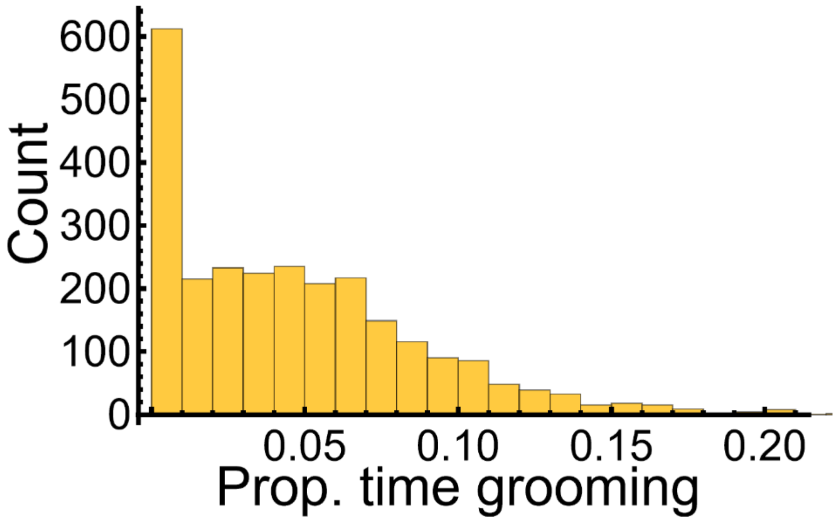


**Figure S3.1**. The raw frequency distribution for the proportion of time adult females spent grooming during focal samples per hydroyear shows zero-inflation in a distribution that is otherwise approximately normal.

**Supplementary Appendix S4: Detailed statistical methods and results for agonistic interaction analyses**

**Table S4.1**. Detailed model results for analyses of agonistic interactions, with response variables listed in italics ($x$ = group size). AIC difference from best-supported model provided for each response variable. P-values associated with the model fit are listed in the third to last column and values associated with fixed effects are bolded when significant (Int = intercept). Response distribution and random effects included in each model are noted in the final two columns. A summary of the best-supported models can be seen in Table 1.

| **Form** | **Model** | **ΔAIC** | **p-values** | **Distribution** | **Random effects** |
|---|---|---|---|---|---|
| | *Aggressive* | | | | |
| Linear | $y = -0.380 + 0.029x$ | 0 | Int:$< 10^{-4}$<br>$\boldsymbol{x}$**:** $\boldsymbol{< 10^{-13}}$ | Gaussian | Group id |
| Linear + quadratic | $y = -0.520 + 0.047x - 0.0005x^2$ | 1.0 | Int: 0.003<br>$\boldsymbol{x}$**: 0.013**<br>$x^2$: 0.332 | Gaussian | Group id |
| | *Non-aggressive* | | | | |
| Linear + quadratic | $y = -0.835 + 0.070x - 0.0009x^2$ | 0 | Int:$< 10^{-8}$<br>$\boldsymbol{x}$**:** $\boldsymbol{< 10^{-6}}$<br>$\boldsymbol{x^2}$**: 0.019** | Gaussian | Group id |
| Linear | $y = -0.580 + 0.038x$ | 3.5 | Int: $< 10^{-12}$<br>$\boldsymbol{x}$**:** $\boldsymbol{< 10^{-15}}$ | Gaussian | Group id |

**Supplementary Appendix S5: Detailed statistical methods and results for analyzing the proportion of positive interactions within dyads**

**Table S5.1**. Detailed model results for the proportion of interactions in a dyad that were positive assuming a binomial distributed error structure ($x$ = group size; $\bar{x}$ = mean group size experienced by a dyad; Int = intercept). The response variables were the number of grooming interactions and number of agonistic interactions for each dyad, represented as ordered pairs. The predictor variables were the difference between the group size a dyad experienced in one hydroyear and the mean group size the dyad experienced across all hydroyears ($x - \bar{x}$; the within-dyad effect) and the mean group size the dyad experienced across all hydroyears ($\bar{x}$.; the between-dyad effect). We also assessed whether the total number of interactions within dyads, $T$, had an effect on whether interactions tended to be positive or negative. Since this quantity is sensitive to observer effort, we used an estimate of $T$ controlled for observer effort as described in Supplementary Appendix S2. Response distribution and random effects included in each model are noted in the final two columns. Significant p-values for coefficients in bold. A summary of the model can be seen in Table 1.

| **Form** | **Model** | **p-values** | **Distribution** | **Random effects** |
|---|---|---|---|---|
| Between-within dyad model | $logit(y) = -0.625 - 0.022(x - \bar{x}) - 0.033\bar{x} + 0.007T$ | Int: $< 10^{-4}$<br>$\boldsymbol{x - \bar{x}}$**:** $\boldsymbol{< 10^{-9}}$<br>$\boldsymbol{\bar{x}}$**:** $\boldsymbol{< 10^{-5}}$<br>$\boldsymbol{T}$**:** $\boldsymbol{< 10^{-15}}$ | Binomial | Group id<br>Dyad id |

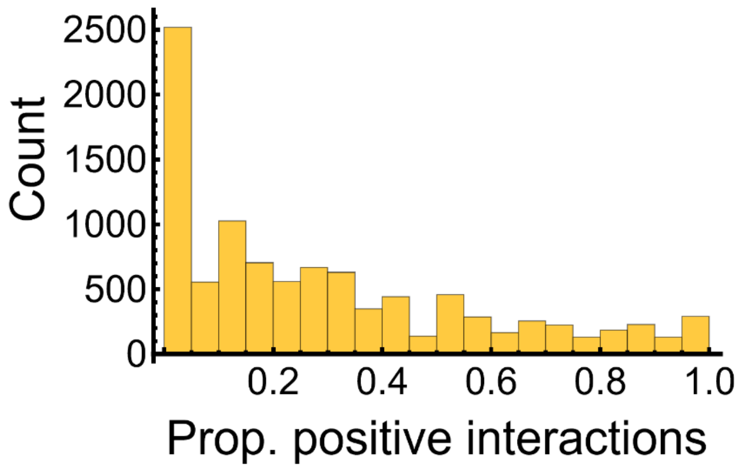


**Figure S5.1**. Raw frequency distribution for the proportion of interactions in a dyad that were positive (i.e., had a positive relationship tenor) across all group-hydroyears. The horizontal axis shows the number of grooming interactions observed for each dyad in a given group-hydroyear, divided by the total number of grooming plus agonistic interactions that they engaged in during the same group-hydroyear.

### Supplementary Appendix S6: Detailed statistical methods and results for social bonding analyses

**Table S6.1**. Detailed model results for mean bond strength with response variables listed in italics ($x$ = group size). AIC difference from best-supported model provided for each response variable. P-values associated with the model fit are listed in the third to last column (Int = intercept). Significant coefficients associated with fixed effects are bolded; data for bond strength and mean weighted degree for "All bonds" were shifted to have a mean of 1, as described in the Methods, to help with model convergence. Response distribution and random effects included in each model are noted in the final two columns. The random effect of group id led to convergence problems for models of shifted mean weighted degree of all bonds and thus was removed. An observation-level random effect ("unique id") was added to some models to lead to better agreement with assumptions. We only present one model for the mean weighted degree of strong bonds due to convergence issues. A summary of the best-supported model for each mean bond strength variable can be seen in Table 1.

| | **Form** | **Model** | **ΔAIC** | **p-values** | **Distribution** | **Random effects** |
|---|---|---|---|---|---|---|
| ***All bonds*** | *Shifted mean bond strength* | | | | | |
| | Linear + quadratic | $\log(y) = 1.936 - 0.134x + 0.003x^2$ | 0 | Int: $< 10^{-15}$<br>***x*: $< 10^{-9}$**<br>***x*²: $< 10^{-7}$** | Gamma | Group id |
| | Linear | $\log(y) = 1.104 - 0.022x$ | 24.7 | Int: $< 10^{-15}$<br>***x*: 0.0001** | Gamma | Group id |
| | *Shifted mean weighted degree* | | | | | |
| | Linear + quadratic | $y = 14.113 + 2.559x - 0.057x^2$ | 0 | Int: 0.005<br>***x*: $< 10^{-4}$**<br>***x*²: 0.003** | Gaussian | N/A |
| | Linear | $y = 27.403 + 0.690x$ | 6.4 | Int: $< 10^{-15}$<br>***x*: $< 10^{-6}$** | Gaussian | N/A |
| ***Only strong bonds*** | *Mean bond strength* | | | | | |
| | Linear + quadratic | $\log(y) = 1.032 - 0.083x + 0.002x^2$ | 0 | Int: $< 10^{-12}$<br>***x*: $< 10^{-5}$**<br>***x*²: 0.001** | Gaussian | Group id<br>Unique id |
| | Linear | $\log(y) = 0.622 - 0.025x$ | 9.3 | Int: $< 10^{-15}$<br>***x*: $< 10^{-11}$** | Gaussian | Group id<br>Unique id |
| | *Mean weighted degree* | | | | | |
| | Linear + quadratic | $\log(y) = 2.507 - 0.087x + 0.002x^2$ | 0 | Int: $< 10^{-15}$<br>***x*: 0.008**<br>***x*²: 0.039** | Gaussian | Group id<br>Unique id |

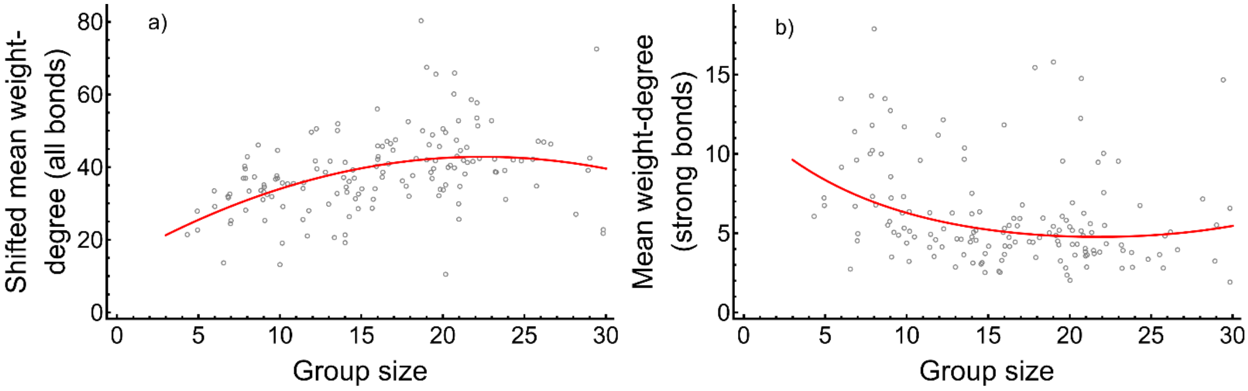


**Figure S6.1**. The mean weighted degree of female-female social bonds was (a) significantly higher in larger groups when considering all bonds, but (b) significantly lower in larger groups when considering only female-female bonds of above-average strength. Circles show mean weighted degree for a year of grooming data for each group and the red lines are the best supported model. As explained in the Methods, data using all bonds were shifted so that the minimum bond strength was one, allowing a log-link function to be used. Thus, the magnitude of the vertical axes are not comparable between panels.

**Table S6.2**. Detailed model results for mean number of social partners as a function of group size, with response variables listed in italics ($x$ = group size). AIC difference from the best-supported model are provided for each response variable. P-values associated with the model fit are listed in the third to last column (Int = intercept) and are bolded when coefficients associated with fixed effects are significant. Response distribution and random effects included in each model are noted in the final two columns. Model assumptions were better met (Fig. S6.3) by including an observation-level random effect for the total number of social partners ("unique id"). A summary of the best-supported model for each response variable can be seen in Table 1.

| | Form | Model | ΔAIC | p-values | Distribution | Random effects |
|---|---|---|---|---|---|---|
| *All bonds* | | *Mean number of total social partners* | | | | |
| | Linear + quadratic | $y = -1.731 + 1.191x - 0.028x^2$ | 0 | Int: 0.199<br>$\boldsymbol{x}$**:** $\mathbf{< 10^{-12}}$<br>$\boldsymbol{x^2}$**:** $\mathbf{< 10^{-8}}$ | Gaussian | Group id<br>Unique id |
| | Linear | $y = 5.325 + 0.241x$ | 30.7 | Int: $\mathbf{< 10^{-13}}$<br>$\boldsymbol{x}$**:** $\mathbf{< 10^{-8}}$ | Gaussian | Group id<br>Unique id |
| *Only strong bonds* | | *Mean number of social partners with bonds of above average strength* | | | | |
| | Linear | $\log(y) = 1.418 + 0.007x$ | 0 | Int: $< 10^{-15}$<br>$x$: 0.251 | Gaussian | Group id |
| | Linear+ quadratic | $\log(y) = 1.545 - 0.010x + 0.0005x^2$ | 1.7 | Int: $< 10^{-10}$<br>$x$: 0.726<br>$x^2$: 0.548 | Gaussian | Group id |

**Supplementary Appendix S7: Detailed statistical methods and results for social network analyses**

**Table S7.1**. Detailed model results for social network metrics ($x$ = group size). AIC difference from best-supported model provided for each response variable. P-values associated with the model fit are listed in the right column (Int = intercept). Significant coefficients associated with fixed effects in bold. Response distribution and random effects included in each model are noted in the final two columns. A summary of the best-supported model for each response variable can be seen in Table 1.

| Form | Model | ΔAIC | p-values | Distribution | Random effects |
|---|---|---|---|---|---|
| | *Clustering coefficient* | | | | |
| Linear + quadratic | $\mathrm{y} = 1.084 - 0.070\mathrm{x} + 0.001\mathrm{x}^2$ | 0 | Int: $< 10^{-15}$<br>$\boldsymbol{x}$**:** $\boldsymbol{< 10^{-9}}$<br>$\boldsymbol{x^2}$**:** $\boldsymbol{< 10^{-5}}$ | Gaussian | Group id |
| Linear | $y = 0.708 - 0.019x$ | 17.6 | Int: $< 10^{-15}$<br>$\boldsymbol{x}$**:** $\boldsymbol{< 10^{-11}}$ | Gaussian | Group id |
| | *Density* | | | | |
| Linear + quadratic | $y = 1.138 - 0.080x + 0.002x^2$ | 0 | Int: $< 10^{-15}$<br>$\boldsymbol{x}$**:** $\boldsymbol{< 10^{-15}}$<br>$\boldsymbol{x^2}$**:** $\boldsymbol{< 10^{-8}}$ | Gaussian | Group id |
| Linear | $y = 0.698 - 0.022x$ | 27.9 | Int: $< 10^{-15}$<br>$\boldsymbol{x}$**:** $\boldsymbol{< 10^{-15}}$ | Gaussian | Group id |
| | *Diameter (subset to connected networks only)* | | | | |
| Linear | $y = 3.706 + 0.055x$ | 0 | Int: $< 10^{-11}$<br>$x$: 0.082 | Gaussian | Group id |
| Linear + quadratic | $y = 2.701 + 0.200x - 0.005x^2$ | 1.2 | Int:0.026<br>$x$: 0.214<br>$x^2$: 0.358 | Gaussian | Group id |

**Table S7.2**. Adding additional controls for observer effort to the best-supported models of social network metrics ($x$ = group size). Now, the response variable is a residual of a social network metric regressed on observer effort (Supplementary Appendix S2). P-values associated with the model fit are listed in the right column (Int = intercept). Significant coefficients associated with fixed effects in bold.

| **Response** | **Model** | **p-values** |
|---|---|---|
| Clustering coefficient | $\mathrm{y} = 0.637 - 0.070\mathrm{x} + 0.002\mathrm{x}^2$ | Int: $< 10^{-10}$<br>$\boldsymbol{x}$**:** $\boldsymbol{< 10^{-8}}$<br>$\boldsymbol{x^2}$**:** $\boldsymbol{< 10^{-5}}$ |
| Density | $y = 0.742 - 0.080x + 0.002x^2$ | Int: $< 10^{-15}$<br>$\boldsymbol{x}$**:** $\boldsymbol{< 10^{-12}}$<br>$\boldsymbol{x^2}$**:** $\boldsymbol{< 10^{-8}}$ |
| Diameter | $y = 0.002 - 0.0001$ | Int: 0.997<br>$x$: 0.997 |
